# Formation of multinucleated globoid cells in Krabbe iPSC-derived microglia cultures: a preliminary investigation

**DOI:** 10.1101/2021.10.16.464657

**Authors:** Richard Lieberman, Grace Gao, Robert B. Hunter, John P. Leonard, Leslie K. Cortes, Robert H. Barker

## Abstract

Globoid cell leukodystrophy (Krabbe disease) is a severe demyelinating, neurodegenerative lysosomal storage disorder caused by deficiency in glycosphingolipid catabolic enzyme galactosylceramidase (GALC). Histologically, Krabbe disease is characterized by the appearance of large multinucleated globoid cells that express classical macrophage markers (both of brain-resident microglia and peripheral monocyte-derived). Globoid cells reside near areas of degeneration; however, their functional significance in disease progression remains unclear. In the current study, we differentiated microglia-like cells from iPSCs from a donor with infantile Krabbe disease and compared them to microglia generated from two healthy controls and two donors with the lysosomal storage disorder metachromatic leukodystrophy (MLD), which is genetically distinct from Krabbe disease but presents similarly in terms of severity of demyelination and neurodegeneration. We report the novel finding of prominent formation of giant multinucleated globoid cells from the microglia derived from the Krabbe donor, but not from healthy control or MLD donors. The Krabbe microglia displayed reduced IL-6 protein expression upon stimulation with lipopolysaccharide, and the multinucleated globoid cells themselves appeared deficient in phagocytosis of both disease-relevant myelin debris and E. coli, together hinting at an impairment of normal function. The formation of the globoid cells could be attenuated by fully replacing the medium following passaging, suggesting that yet-to-be determined secreted factors are influencing cell fusion in our culture system. While preliminary, our results imply that globoid cells may be detrimental in Krabbe disease by hindering the normal function of brain-residing macrophages.

## Introduction

Sphingolipids, including glycosphingolipids, comprise a complex set of sphingoid base-containing lipids that have diverse biological roles (1). Glycosphingolipids are highly expressed in the brain and are major components of neural membranes, contributing to cellular structure, direct cell-to-cell interaction, and intra- and inter-cellular signaling, among other functions including those continuing to be discovered (2-4). Dysregulation of glycosphingolipid metabolism caused by mutations in genes encoding lysosomal enzymes can lead to accumulation of undegraded substrate and subsequent neuropathology, in severe cases manifesting within the first years of life (5, 6). While many of the genetic defects that result in lysosomal storage disorders are known, the downstream cellular cascades that influence neurodegeneration remain to be fully elucidated.

Globoid cell leukodystrophy (Krabbe disease), a rare (∼1:100,000 births) lysosomal storage disorder hallmarked by the appearance of multinucleated globoid cells in the brain, is characterized by progressive neurodegeneration and demyelination of the central, and to a lesser extent, peripheral nervous systems (7, 8). The disease is classified by age of onset with the most severe infantile-onset form accounting for 85-90% of cases, which typically manifests prior to 12 months of age and has an average age of death around 24 months (9). Krabbe disease is caused by deficiency of the enzyme galactosylceramidase (GALC) arising from recessive, deleterious mutations in its gene, *GALC* (10-12). GALC deficiency results in the inability to degrade galactosylceramide (Figure 1), a glycosphingolipid highly expressed in myelinating cells of the nervous system (13), leading to a paradoxical decrease in galactosylceramide in postmortem Krabbe nervous system tissue but an increase in its toxic metabolite galactosylsphingosine (psychosine) compared to healthy controls (14). This is believed to be due to the protective effect of GALC hydrolysis of psychosine under normal conditions (14). These findings, in tandem with results highlighting that psychosine exposure has effects on cells of the brain (15) including oligodendrocytes (16, 17), neurons (18, 19), astrocytes (20, 21), and microglia (22, 23), has led to the acceptance of the “psychosine hypothesis,” which postulates that psychosine is the main driver of disease progression (14, 24).

**Figure 1.**
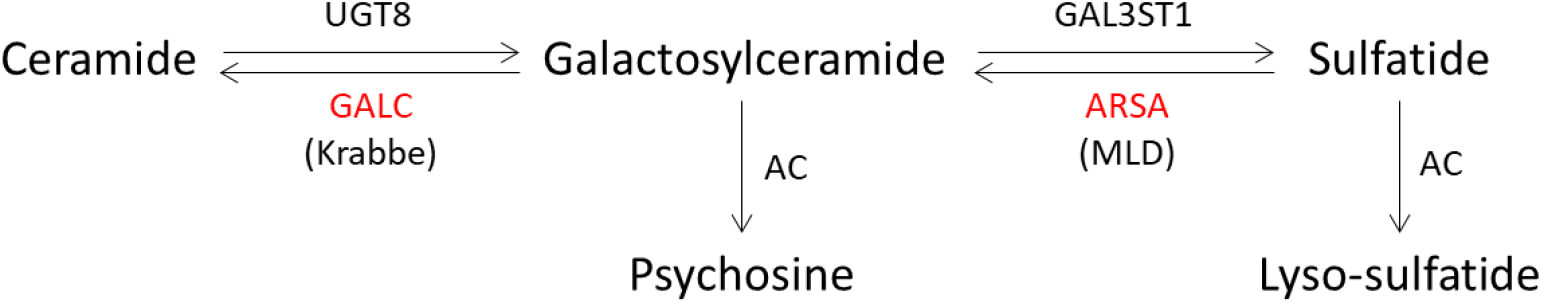
Schematic of the biosynthesis of glycosphingolipids relevant to Krabbe disease and MLD. A simplified version of the lipid biosynthesis pathway depicts the relationship between Krabbe disease and MLD. Krabbe disease is caused by deficiency in GALC leading to the impaired catabolism of galactosylceramide. Galactosylceramide is deacylated by acid ceramidase (AC) to produce psychosine, which is currently believed to accumulate to toxic levels in Krabbe disease. Just downstream of galactosylceramide is sulfatide, which is catabolized via the actions of arylsulfatase A (ARSA). Deficiency of this enzyme causes MLD and results in the accumulation of sulfatide and its deacylated form lyso-sulfatide.

The namesake multinucleated globoid cells that form in the brains of Krabbe patients appear around areas of demyelination (25, 26). Multinucleated globoid cells express classical macrophage markers including IBA1 and CD11b, and in the disease state arise from cellular fusion primarily of brain-resident microglia but also from infiltrating peripheral macrophages (23, 27). Their relevance in Krabbe disease pathology remains to be fully elucidated. Globoid cell occurrence is inversely correlated with Krabbe patient longevity (26), suggesting that they are either detrimental to disease progression and/or form in response to detrimental drivers of disease, for example psychosine (22, 23). Since giant multinucleated cells form *in vitro* from exogenous stimulation with inflammatory factors and *in vivo* from inflammation arising from bioimplants and prosthetics (28-30), it has been suggested that the neuroinflammation caused by demyelination results in the formation of globoid cells in Krabbe disease (31). However, globoid cells are observed in a murine model of Krabbe disease in the absence of demyelination and neuroinflammation (32). These conflicting results highlight the need for more information to fully address the role of globoid cells in the pathogenesis of Krabbe disease.

Metachromatic leukodystrophy (MLD) is a rare (∼1:60,000 births) lysosomal storage disorder that arises mainly due to deficiency in the lysosomal enzyme arylsulfatase A, caused by deleterious recessive mutations in its gene, *ARSA* (33, 34). Arylsulfatase A is adjacent to GALC in the galactosylceramide metabolic pathway and catabolizes sulfatide into galactosylceramide (35) (Figure 1). Deficiency in the enzymatic activity of arylsulfatase A leads to accumulation and lysosomal storage of its substrate sulfatide and lyso-sulfatide, that subsequently result in cellular dysfunction (36, 37). The clinical manifestation of MLD is similar to Krabbe disease, including infantile demyelination and neurodegeneration in the most severe form, which is a reason why Krabbe disease and MLD are considered the two classic genetic leukodystrophies (7, 10). Aberrant microglial function is common between the two diseases (31, 38, 39), and hematopoietic stem cell replacement therapy is somewhat therapeutically efficacious in the course of both diseases (40-42), suggesting that, at least in part, cells of this lineage are important to disease progression. However, histologically Krabbe disease is distinct from MLD due to the prominent formation of multinucleated globoid cells in the brain (43). This implies that, while both diseases have common microglial abnormalities, factors distinct to the pathology of Krabbe disease result in a different microglial phenotype than in MLD.

A human cell model may provide valuable insights into the role of microglia in Krabbe disease and could be useful in identifying differences between Krabbe and MLD microglia that result in the Krabbe-specific globoid cells. Pluripotent stem cells generated from human fibroblasts (44) have the capacity to differentiate into many lineages of the central nervous system (45). Recent protocols describe the generation of phagocytically-competent iPSC-derived microglia-like cells that are similar to fetal human microglia at the transcriptomic level (46-50), suggesting that human iPSCs provide a tool to explore microglial pathology in Krabbe disease. Therefore, we generated microglia-like cells from a donor with Krabbe disease and compared them to cells generated from two healthy controls and two donors with MLD. Unexpectedly, but in line with disease characteristics, we identified robust formation of multinucleated globoid cells in microglia cultures derived from the Krabbe donor, recapitulating the hallmark of the disease. Globoid cell formation was observed in the absence of exogenous psychosine administration, which has been used in prior studies to induce multinucleated cells (22, 23). We used this phenotype to qualitatively probe the consequences of cell fusion in our novel cell model utilizing assays relevant to microglial function. Our findings are in support of prior studies suggesting an important role of microglia in the pathogenesis of Krabbe disease (for review, see (31)).

## Materials and Methods

### iPSC generation

Fibroblasts from two healthy donors (Control 1: GM05659, 1 year old male; Control 2: HUCMF05, male cord blood), two donors with infantile onset MLD (MLD1: GM00197, 4 year old male; MLD2: GM00905, 3 year old female), and one donor with infantile onset Krabbe disease (GM06806, 2 year old female) were obtained from the Coriell Institute for Medical Research, with the exception of Control 2, which was obtained from Icelltis. Reprogramming to pluripotency was carried out using the CytoTune 2.0 sendai virus kit (Thermo-Fisher Scientific) to overexpress four transcription factors (OCT4, SOX2, KLF4, and c-Myc) according to the manufacturer’s protocol. Individual iPSC colonies derived from each donor were manually selected and clonally expanded on Matrigel (Corning) coated plates in complete mTeSR1 medium (StemCell Technologies). Cultures were monitored and areas of spontaneous differentiation were removed. Cells were passaged when they reached ∼80-90% confluence using ReLeSR (StemCell Technologies). Induction of pluripotency was validated by the expression of OCT4 and NANOG via immunofluorescence (data not shown). G-banded karyotyping was carried out by the WiCell Institute (Madison, WI). All iPSC lines displayed a normal karyotype except for MLD2, which was found to have a duplication of the long arm of chromosome 14 in 7 of 20 cells examined. The line was included in the current study due to its appropriate morphology, expression of pluripotent markers, and its ability to differentiate robustly and successfully.

### Differentiation of microglia-like cells

Microglia-like cells were differentiated utilizing a serum-free protocol containing aspects of three published protocols (49-51). In sum, this protocol is designed for the continuous harvesting of primitive macrophage precursors that are shed from embryoid bodies over ∼6-8 months for terminal differentiation into microglia using defined factors. First, embryoid bodies (EBs) were generated from confluent iPSC cultures by dissociating them to single cells and seeding them at a density of ∼4 × 10^6^ cells per well of Aggrewell 800 6-well plates (StemCell Technologies), equating to ∼2200 cells per EB. Cells were seeded and cultured in EB medium consisting of complete mTeSR1 supplemented with 10 µM Y27632 (StemCell Technologies), 50 ng/mL BMP-4, 20 ng/mL stem cell factor (SCF), and 50 ng/mL VEGF-121 (all from Peprotech). EB medium was replaced every other day for 7 days. EBs were collected from the Aggrewell plates, centrifuged and washed in X-VIVO 15 medium (Lonza Biosciences) and seeded into tissue culture treated vessels at a density of 16 EBs per well of a 6-well plate, 75 EBs per T75 flask, or 150 EBs per T150 flask. Cells were seeded and cultured in hematopoietic cell medium consisting of X-VIVO 15 medium (with gentamicin and phenol red) supplemented with 2mM Glutamax (Gibco), 1x penicillin/streptomycin (Gibco), 55 µM β-mercaptoethanol (Gibco), 100 ng/mL M-CSF (Peprotech), and 25 ng/mL IL-3 (Cell Guidance Systems). Following seeding of EBs, cultures were not agitated for 7 days to allow for adherence. Lines with poor adherence efficiency could be seeded at higher densities for better downstream yield (we have used up to 50 EBs in a well of a 6-well plate). An equal volume of fresh hematopoietic cell medium was added to the cultures on days 7 and 14 following seeding, with no medium being removed. Thereafter, ∼75% of the medium was collected and replaced with fresh hematopoietic cell medium every 7 days.

Primitive macrophage precursors (PMPs) could be observed in suspension shedding from the adhered EBs for up to ∼6-8 months in culture. The PMPs were harvested during the medium change by collecting supernatant through a 40 µm filter to remove unwanted debris. Cells were centrifuged and resuspended in microglia medium consisting of Advanced DMEM/F12 (Gibco) supplemented with 2mM Glutamax. 1x penicillin/streptomycin, 1x N2 supplement (Gibco), 100 ng/mL IL-34, and 10ng/mL GM-CSF (both from Peprotech). 2.5 million or 7.5 million cells were seeded into tissue culture treated 10 CM to 15 CM plates, respectively. Fresh microglia medium was fully replaced every 2-3 days for 7 days.

Following terminal differentiation microglia were removed from the plate via a 10-minute incubation with Accumax (Millipore) and lifted with a cell scraper. Cells were collected, centrifuged, and resuspended in complete microglia medium for counting. The passaged microglia were directly seeded in complete microglia medium into assay plates. In one experiment, 1 nM or 10 nM rhGALC protein (R&D Systems) was added to the microglia medium at the time of seeding into the assay plate.

### Live cell imaging with Calcein Green AM

Differentiated microglia were seeded into tissue culture treated 96-well high content imaging plates (CellCarrier Ultra, Perkin Elmer) at a density of 60,000 cells/well. 96 hours after seeding medium was aspirated and replaced with PBS containing calcium and magnesium supplemented with Hoechst 33342 (1:1000, Thermo-Fisher Scientific) and Calcein Green AM (50 µg vial resuspended in 50 µL DMSO, 1:1000, Thermo-Fisher Scientific) to identify live cells. Cells were incubated with dye for 10 minutes at 37C and then directly imaged using a 5x objective on a Perkin Elmer Opera Phenix high content microscope.

### Phagocytosis of pHrodo conjugated substrates

Phagocytosis of pHrodo-conjugated *E. coli* and myelin debris was performed utilizing published methods (52, 53). pHrodo Red-labeled *E. coli* (Thermo Fisher Scientific) was resuspended in PBS to 1 mg/mL, vortexed vigorously, sonicated for 5 minutes, and used fresh or stored at 4°C until use, according to the manufacturer’s instructions. Myelin debris was separated via Percoll (Sigma Aldrich) gradient, pelleted, and washed three times with PBS. Protein concentration was determined via BCA and 4 mg/mL aliquots in PBS were stored at -80C until use. Labeling of myelin debris with pHrodo Red SE (Thermo-Fisher Scientific) was carried out according to the manufacturer’s instructions. First, 7.5% (0.9M) sodium bicarbonate buffer was added to ∼800 µg of myelin debris in PBS to a final concentration of 100 mM. pHrodo Red SE was resuspended to 10 mM in DMSO and 1.5 µL was added to the 800 µg myelin solution and incubated at room temperature for 45 minutes. Conjugated myelin was pelleted via centrifugation at 12,000 RPM for 5 min and washed three times to remove unbound pHrodo dye. Supernatant from the final wash was collected and stored at -80°C and used as a loading control. pHrodo-conjugated myelin was resuspended to 500 µg/mL in PBS and stored at -80°C until use.

Healthy control and Krabbe microglia were passaged into 96 well high content imaging plates at a density of 60,000 cells/well and maintained in culture for 96 hours to allow for the formation of multinucleated cells prior to the phagocytosis assay. pHrodo-conjugated substrate was thawed and resuspended to 20 µg/mL in complete microglia medium described above. Medium was aspirated from the microglia and replaced with the medium supplemented with substrate and incubated at 37°C for 2 hours. Prior to live cell imaging medium was aspirated and replaced with Hoechst (1:1000)-containing PBS (+calcium and magnesium) and cells were incubated for 5 minutes at 37°C. Images were acquired using a 20x objective on a Perkin Elmer Opera Phenix microscope.

### Measurement of IL-6 expression

Microglia from two healthy controls, 2 MLD donors, and the Krabbe donor were differentiated for 7 days and passaged into a 96-well plate at 60,000 cells/well. Cells were cultured for 96 hours and then stimulated with medium (mock condition) or 50 ng/mL lipopolysaccharide (LPS, purchased from Sigma-Aldrich) by spiking in 500 ng/mL LPS at a 1:10 ratio (i.e. 20 µL of 500 ng/mL LPS into the 200 µL culture medium). Cells were incubated with LPS for 24 hours. Supernatant was collected, diluted 1:5 (mock condition) or 1:20 (LPS condition) and analyzed on a PHERAstar plate reader (BMG Labtech) using alphALISA technology (Perkin Elmer) per the manufacturer’s instructions. Concentration of IL-6 was calculated based on the standard curve provided in the alphALISA kit. Following collection of the supernatant, cells were live imaged with Calcein Green AM and Hoechst, as described above, and the entire well was imaged at 5x magnification. The number of viable nuclei was quantified by counting the number of Calcein positive nuclei using Harmony software (Perkin Elmer). Concentration of IL-6 protein was then normalized to the total viable nuclei count to generate concentration on a per-viable nuclei basis, which allowed for comparison between wells and donors and controlled for globoid cell formation and discrepancies in seeding density. 8 wells total of a 96-well plate were measured per donor subject, 4 wells for baseline, and 4 LPS-stimulated. One-way ANOVA with post-hoc Dunnett’s multiple comparison test was used for statistical analysis in GraphPad Prism Version 8 for Windows (GraphPad Software, www.graphpad.com).

### Immunocytochemistry

Cells were fixed in 4% paraformaldehyde for 15 minutes at room temperature, washed twice with PBS, and blocked and permeabilized in 10% normal donkey serum (Abcam) supplemented with 0.3% triton X-100 (Sigma-Aldrich) for 15 minutes at room temperature. Blocking solution was aspirated and cells were washed with PBS. Primary antibodies were added in antibody dilution buffer consisting of 2% normal donkey serum, 1% bovine serum albumin (Sigma-Aldrich) and 0.1% TWEEN-20 (Sigma-Aldrich) in PBS. The following primary antibodies were used: mouse anti-OCT4 (1:1,000; Millipore MAB4419), goat anti-IBA1 (1:500; Abcam Ab5076), and rabbit anti-CD11b (1:500, Abcam Ab133357). Primary antibodies were incubated overnight at 4°C, following which cells were washed four times in PBS and donkey anti-mouse 488, donkey anti-rabbit 488, or donkey anti-goat 568 Alexa Fluor-conjugated secondary antibodies (diluted 1:1,000 in antibody dilution buffer; Thermo-Fisher Scientific) were applied at room temperature for 2 hours. Cells were washed four times with PBS and CellMask Deep Red (1:10,000; Thermo-Fisher Scientific) and Hoechst 33342 (1:10,000; Thermo-Fisher Scientific) diluted in PBS were applied to the cells for 10 minutes at room temperature, following which cells were washed two times with PBS and imaged on a Perkin Elmer Opera Phenix microscope.

## Results

### Formation of large multinucleated globoid cells in Krabbe microglia-like cells

Microglia-like cells were differentiated from iPSCs derived from 2 healthy controls, 2 donors with MLD, and 1 donor with Krabbe disease (Figure 2A-E). Following 7 days in microglia differentiation medium, cells were enzymatically and mechanically passaged into microwell assay plates. We used the viability probe Calcein Green AM to examine the gross morphology of the cultures. 4 days after passage we noted sporadic cell fusion in healthy control and MLD-derived cultures (Figure 2F, white arrows, Supplementary Figure 1). The formation of fused, multinucleated globoid cells was prominent and robust in cultures derived from the Krabbe donor, which appeared to be larger and more pronounced than the sparse multinucleated cells present in healthy control and MLD-derived cultures (Figure 2F, red arrow, Supplementary Figure 1).

**Figure 2.**
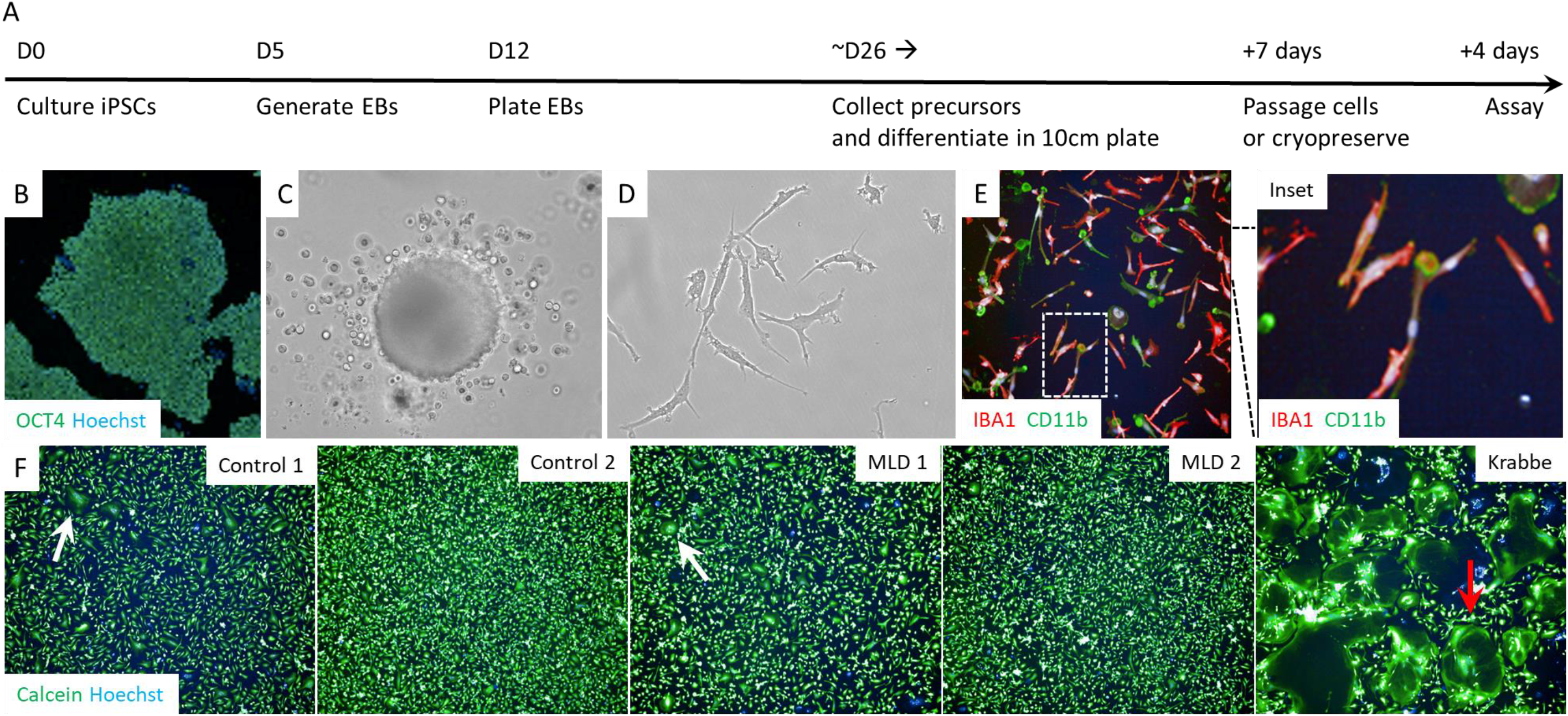
Formation of multinucleated globoid cells in microglia derived from Krabbe iPSCs. (A-E) Timeline and representative images of the differentiation of microglia-like cells. OCT4-expressing pluripotent iPSC colonies (B) were seeded into Aggrewell plates to generate EBs, which were cultured in medium facilitating the continuous shedding of primitive macrophage precursors (C). Precursors were collected from the medium and plated in defied medium for terminal differentiation into cultures of microglia-like cells (D) expressing IBA1 and CD11b (E). (F) After 7 days in microglia medium cells were lifted from the culture vessel and seeded into assay-format microwell plates. 4 days after passaging, microglia were imaged with the live cell fluorescent probe Calcein Green AM. Cell fusion was observed during this time in all cultures. Multinucleated cells formed in healthy control and MLD cultures (white arrows). The formation of multinucleated globoid cells was more robust in microglia cultures derived from the Krabbe donor (red arrow). Images in panels B-E were obtained with a 20x objective, and all images in panel F were obtained with a 5x objective.

### Impaired microglial function observed in Krabbe cells

We examined IL-6 production following LPS stimulation and phagocytosis of fluorophore-conjugated substrates to assess the functional significance of globoid cell formation. Microglia were stimulated with 50nM LPS following globoid cell-induction by passaging, which resulted in robust expression of IL-6 protein in all cultures. There was significantly more IL-6 protein (Figure 3) measured in the supernatant of control and MLD cultures compared to the Krabbe cells (ANOVA: F (4, 15)= 6.96, p = 0.002; Dunnett’s multiple comparisons post-hoc test: Krabbe vs. Control 1; p = 0.002, Krabbe vs Control 2; p = 0.026, Krabbe vs. MLD1; P = 0.001, Krabbe vs MLD2; p = 0.009).

**Figure 3.**
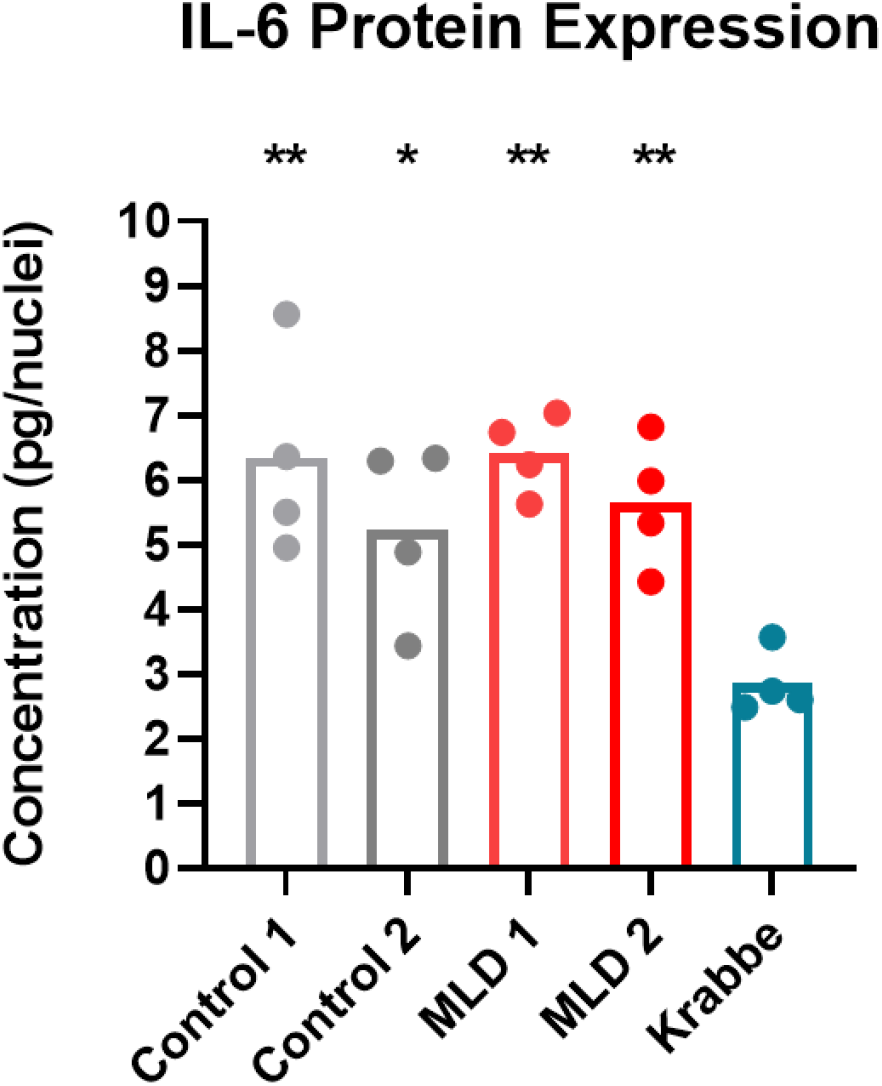
Krabbe microglia have reduced production of IL-6 following LPS stimulation. Microglia derived from the two healthy controls, 2 MLD donors, and a donor with Krabbe disease were passaged and seeded into assay-format 96 well plates, cultured for 96 hours, and then stimulated with 50 ng/mL lipopolysaccharide (LPS) for 24 hours. IL-6 protein was measured in the supernatant of the cultures by alphALISA and expression was normalized to the total viable nuclei count obtained via imaging of the assay plate. Globoid-cell containing Krabbe cultures displayed reduced LPS-stimulated IL-6 protein levels compared to the microglia generated from other subjects. Basal IL-6 was not detected in any of the cultures. Each dot represents the protein expression data obtained from a single well of a 96 well plate. * p < 0.05, ** p < 0.01

Microglia from one healthy control and the Krabbe donor were passaged into assay plates to induce globoid cell formation. Cultures were then incubated with pHrodo-conjugated substrates, either mouse myelin debris or *E. coli*. The pHrodo probe is only fluorescent when internalized and localized to an acidic environment. Following a 2-hour incubation with substrate, we observed a lack of fluorescent signal of conjugated myelin (Figure 4A; Supplementary figure 2) and *E. coli* (Figure 4B) in multinucleated globoid cells. Importantly, non-multinucleated microglia derived from the Krabbe donor displayed functional phagocytosis of these substrates, suggesting that they are phagocytically competent when not fused.

**Figure 4.**
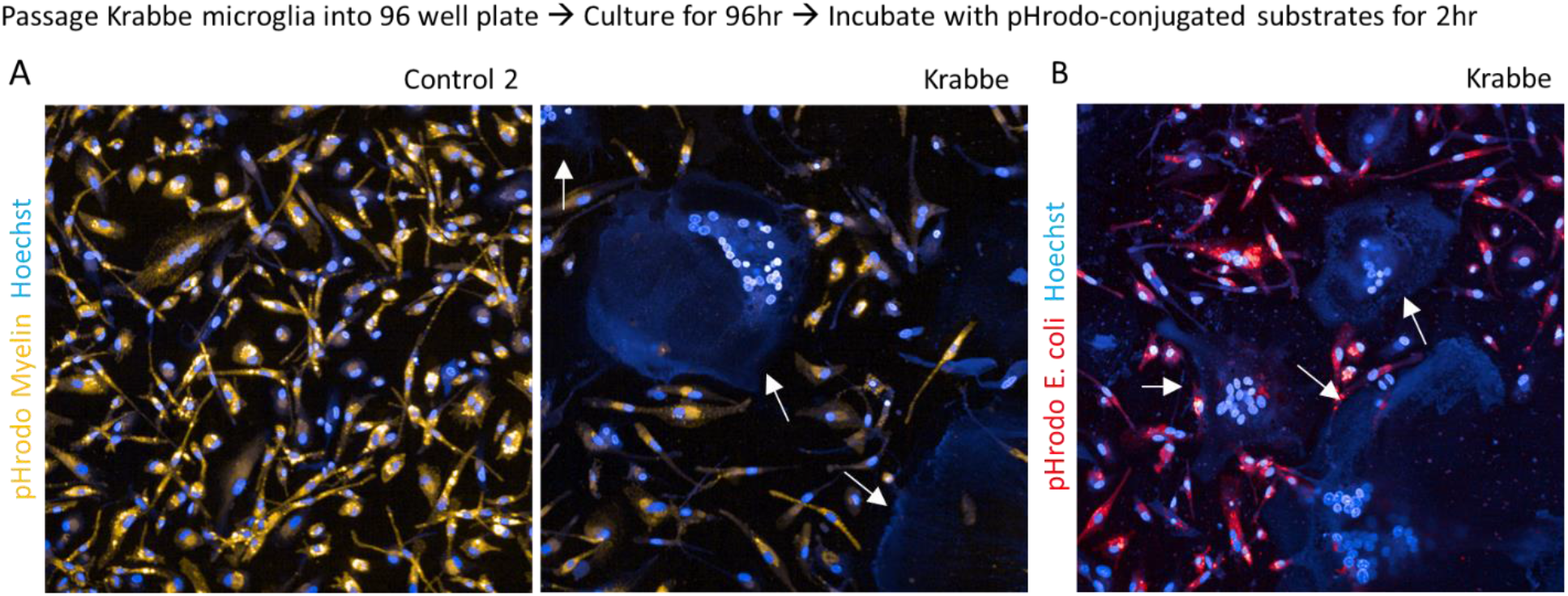
Impaired phagocytosis of myelin and E. coli in multinucleated globoid cells. Differentiated microglia derived from a healthy control and Krabbe donor were passaged and seeded into assay-format 96 well plates, cultured for 96 hours, and then incubated with pHrodo-conjugated substrates for 2 hours. Fluorescence of pHrodo-conjugated myelin debris (A) and *E. coli* (B), indicating its internalization and acidic-vesicle localization was absent in multinucleated globoid cells. Phagocytosis of substrates was apparent in individual microglia in the control and Krabbe cultures. White arrows denote globoid cells lacking fluorescence of conjugated substrate. Images were obtained with a 20x objective. pHrodo myelin is pseudo colored orange, and pHrodo *E. coli* is pseudo colored red.

### Globoid cell formation can be attenuated by medium replacement

Extracellular matrix proteins modulate the formation of globoid cells and alter microglial functions in models of Krabbe disease (22, 23), and culture medium supplemented with cytokines including IL-4 or IL-13 induces the formation of giant multinucleated cells in bone marrow derived macrophages (28, 29,54). With respect to these published findings, we explored if factors secreted into the medium following passaging might be influencing cell fusion. As a pilot experiment, we induced globoid cell formation by passaging Krabbe microglia and fully replaced the medium at 2-, 4-, and 24-hours post-plating. Cells were left in culture until the 96-hour time point was reached. In comparison to a condition with no medium change, we noted a visible reduction in globoid cell formation in wells where the medium had been replaced at 2- and 4-hours post-plating. The attenuation of globoid cell formation was not noticeable in wells where the medium had been replaced 24-hours after seeding (Figure 5).

**Figure 5.**
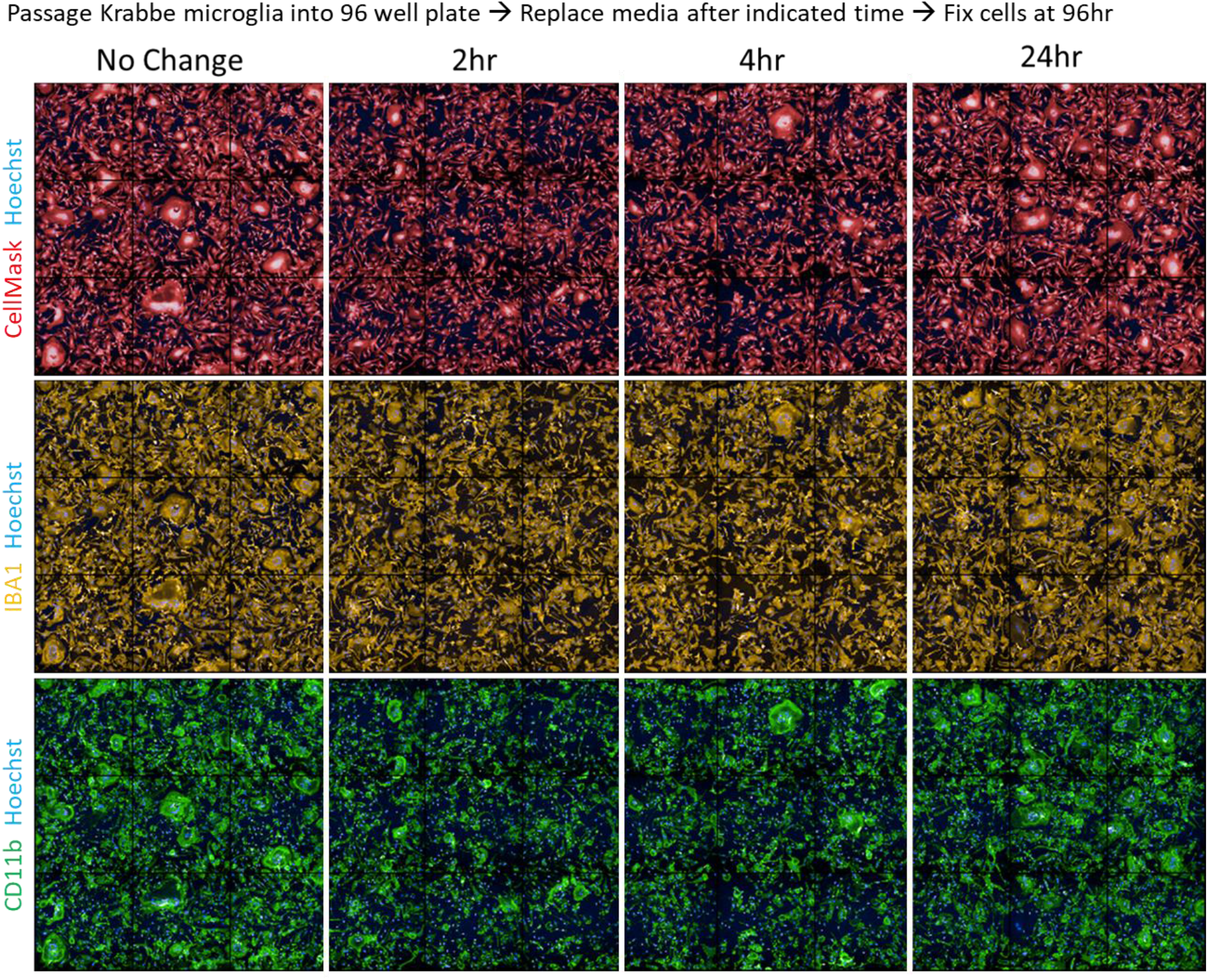
Attenuation of globoid cell formation in Krabbe microglia by medium replacement following passaging. Microglia from the Krabbe donor were passaged and seeded into a 96 well assay plate. Cells adhered to the tissue culture-treated plate within minutes following passaging. 100% of the medium was aspirated and replaced with fresh microglia medium at the indicated times. Cells were cultured for 96 hours prior to fixation. We observed that the formation of globoid cells was attenuated in wells where the medium was replaced 2 hours and 4 hours following seeding. Globoid cells expressed macrophage markers IBA1 (orange) and CD11b (green). Images were obtained with a 20x objective and depict 9 adjacent fields.

## Discussion

iPSCs can be differentiated into disease-relevant cell types that may facilitate the identification of phenotypes related to disease pathogenesis. In the current study, we capitalized on published methods (49-51) to differentiate IBA1+ microglia-like cells from a donor with Krabbe disease and compared them to cells generated from two heathy donors and two donors with MLD. To our knowledge, this proof-of-concept report is the first to demonstrate the ability of iPSC-derived microglia to be generated from patients with Krabbe disease and MLD, suggesting that researchers can take advantage of this workflow to probe the disease biology of this cell type *in vitro*.

We opted to follow the differentiation protocols established by Brownjohn el al., 2018 and Haenseler et al., 2017 because of the detailed characterization of the cells in these publications. Using these culture methods, iPSCs generate nearly pure cultures of microglia-like cells that express classical markers IBA1, CD45, and TREM2, are phagocytically competent, and upregulate cytokines following LPS stimulation (49, 50). Transcriptomic profiling identified that microglia differentiated using these methods clustered with primary human microglia cultured *in vitro*, although it is noteworthy that the iPSC microglia clustered distinctly from profiles of human microglia and macrophages *ex vivo* (49). However, Haenseler et al., 2017 found that co-culture of iPSC microglia with iPSC-derived mixed neuron cultures lead to transcriptomic clustering with fetal human microglia (50), perhaps suggesting that cell signaling factors or direct cell interactions are necessary and sufficient to push iPSC-derived microglia to a more mature state that resembles the normal human developmental state. Indeed, Takata et al. 2017 also found that co-culture of iPSC-derived macrophage precursors with neurons was necessary to drive the cells to an embryonic microglia-like state (55). Taken together, these results suggest that the methods used in our current study to generate iPSC-derived microglia are sufficient to probe the biology of Krabbe disease and other lysosomal storage disorders characterized by microglia involvement, but results should be considered with respect to the overall immaturity of the cells and the lack of co-culturing with neurons.

Robust formation of large multinucleated globoid cells was observed in microglia generated from the donor with Krabbe disease, recapitulating a histological hallmark of the disease. The formation of these cells was more prominent in the Krabbe-derived cultures than either healthy control-derived or MLD patient-derived cultures. Using two healthy donors and two donors with MLD, a genetically distinct demyelinating disease that manifests similarly to Krabbe disease but lacks the formation of globoid cells (7), allowed us to be more confident in the phenotype identified in the Krabbe line. We probed the functional significance of globoid cell formation in the Krabbe microglia with pHrodo-labeled myelin debris or *E. coli* to visually identify phagocytically-competent cells. Strikingly, the globoid cells were deficient in phagocytosis, indicated by the lack of fluorescence signal in the multinucleated cells. This finding best complements recent evidence in murine models that heterozygous mutations in the *GALC* gene resulted in microglial impairments of myelin clearance *in vivo* and *in vitro* (52). Furthermore, macrophage depletion in the mouse model of Krabbe disease (the *twitcher* mouse) resulted in increased myelin debris and exacerbated disease progression (56). Impaired clearance of debris by microglia has been shown to increase with aging, causing cellular senescence and dysfunctional immune responses, and is implicated in the pathogenesis of other neurodegenerative diseases including multiple sclerosis and Alzheimer’s (57, 58), suggesting that proper phagocytic function is an important aspect of neural homeostasis. With respect to our observation of deficient phagocytosis in the human Krabbe globoid cells, these findings further support the observation that impairment of microglial phagocytosis may contribute to Krabbe disease pathology.

Cellular function was also probed by stimulating all of the cell lines with LPS and measuring protein expression of the pro-inflammatory cytokine IL-6, which was selected as a readout because it is upregulated in Krabbe patients and the *twitcher* mouse and implicated in the cellular response to psychosine (59-62). While expression was not detected in the basal state, LPS stimulation resulted in the upregulation of IL-6 in all cultures, validating their differentiation into an LPS-responsive cell type. Interestingly, the Krabbe cultures had markedly reduced IL-6 expression compared to the healthy control and MLD cells. This result complements the finding that IL-6 deficiency exacerbated disease progression in the *twitcher* mouse, including increased number of PAS positive cells, reduced time to the twitching phenotype, and exaggerated gliotic response (63), and suggests that in the disease state IL-6 may have a protective function (61) that is impeded by impaired microglial function. However, our result is not in agreement with the overall elevated IL-6 observed in Krabbe patient brain, but it is worth noting that this inflammatory cytokine arises from other cell sources in Krabbe disease, including astrocytes (20, 59, 60). A limitation of our current study was the examination of a single inflammatory cytokine, when multiple have been implicated in disease (61). Future studies can probe dysregulated expression of additional cytokines in the iPSC derived Krabbe microglia using a cytokine profiling panel, which might offer novel insights into the neuroimmune aspects of the disease. Together, our findings from both the phagocytosis assay and LPS-stimulation experiment suggest an impairment of normal microglial function of the Krabbe microglia cultures.

It is important to highlight that our finding of impaired phagocytosis of myelin and *E. coli* in globoid cells is in contrast with *in vitro* evidence in psychosine-induced globoid cells generated from mouse microglia, which show an increase of phagocytosis of fluorescently labeled beads (23). One possible explanation for this discrepancy is that globoid cells might resemble giant multinucleated cells, also called foreign body giant cells, which form from the fusion of macrophages in response to inflammation following implants of biomaterials, prosthetics, or medical devices (30). These cells function to eliminate foreign material that is too large for individual macrophages (30, 64). Therefore, it is possible that multinucleated globoid cells are primed to phagocytose larger objects, such as beads, which may not be as physiologically relevant, rather than disease relevant substrates such as myelin or dead cells. In support of our speculation of the similarity between the cell types, data suggests that foreign body giant cells display an attenuated pro-inflammatory cytokine profile (65), which complements our finding of reduced IL-6 expression in the Krabbe derived globoid cell-containing cultures following LPS treatment. Future studies could examine differences in selectivity of substrate internalization in macrophages and fused globoid cells by comparing phagocytosis of myelin and conjugated beads in the described human iPSC globoid cell model.

Globoid cell formation appeared to be attenuated by changing the medium within a short window following passaging, suggesting that factors secreted from the cells are influencing the formation of multinucleated cells and that globoid cell formation can be modulated in the presence of the disease-causing *GALC* mutation. Of note, we were not able to attenuate globoid cell formation in an experiment where 1 nM or 10 nM recombinant human GALC protein was supplemented into the medium at the time of seeding (Supplementary Figure 3). This approach was limited however, as we are not able to determine the amount of functional GALC protein entering the cell or confirm its correct localization to the lysosome and effect on lysosomal glycosphingolipids. Considering previous work examining factors and downstream pathways influencing globoid cell formation in non-Krabbe derived cells, including the cytokines IL-4 and IL-13 (28, 54, 64), we hypothesize that 1) mutations in *GALC* render our iPSC-derived microglia-like cells into a primed state to fuse, and that the stress of passaging and seeding releases factors that trigger fusion, or 2) that the factors influencing cell fusion in our culture system are modulated to a greater degree following passaging in Krabbe-derived microglia than in the control or MLD cultures, leading to the increased fusion observed in the Krabbe patient cells. We note that these hypotheses do not have to be mutually exclusive, and further knowledge on the structure and function of the GALC protein under normal physiologic conditions (66, 67) will provide valuable insight towards addressing these questions. A limitation of our study is that we did not evaluate the effect of media change on the smaller fused cells that appeared in control and MLD microglia cultures, as our primary focus was on the globoid cells that formed in the Krabbe cultures due to the large qualitative window provided with our cellular imaging. Therefore, we do not know if the timing of media change would have a similar effect of reducing cell fusion in those cell cultures. Future studies can utilize an iPSC-approach to address if mechanisms leading to cell fusion are conserved across healthy and disease cell lines.

The overall limitations of our pilot investigation need to be considered when assessing its findings. First, we were limited to the use of a single Krabbe-derived iPSC line. Although we attempted to account for this limitation by utilizing two MLD-derived lines in addition to two healthy control lines for comparison, additional iPSC-derived microglia generated from multiple donors, including lines derived from the severe infantile onset and milder late onset forms, will better inform researchers on the role of microglia in disease progression and globoid cell formation. Next, we did not examine substrate accumulation in the Krabbe and MLD microglia and therefore do not know if the microglia-like cells used in the current study recapitulate lipid accumulation observed in the disease state. To this end, we were unable to determine if perturbed lipid levels affected globoid cell formation, which should be addressed in future studies. Finally, we did not probe the factors that might be influencing globoid cell formation in our culture system. There are multiple proteins, both soluble and membrane-bound, that regulate the formation of giant multinucleated cells (68, 69). We speculate that iPSC-derived microglia will be a useful model to identify mediators of cell fusion relevant to Krabbe disease.

In conclusion, we demonstrate the feasibility of a human iPSC model to explore microglial function in Krabbe disease and MLD, which can be extended to examine other lysosomal storage disorders where microglia are of interest. The novel finding that iPSC microglia recapitulate the hallmark phenotype of globoid cell formation in Krabbe disease allowed us to probe the biology of these cells, whose functional significance in disease progression remains incompletely understood. Results from our study largely corroborate prior findings and offer additional insight into Krabbe disease biology in relevant human cells *in vitro*. As globoid cell formation has been observed in other neurologic diseases including amyotrophic lateral sclerosis (70), HIV encephalitis (71), and Alzheimer’s (72), we postulate that iPSC microglia will serve as valuable models to explore commonalities that might be shared across multiple diseases.

---

**Supplementary figure 1.**
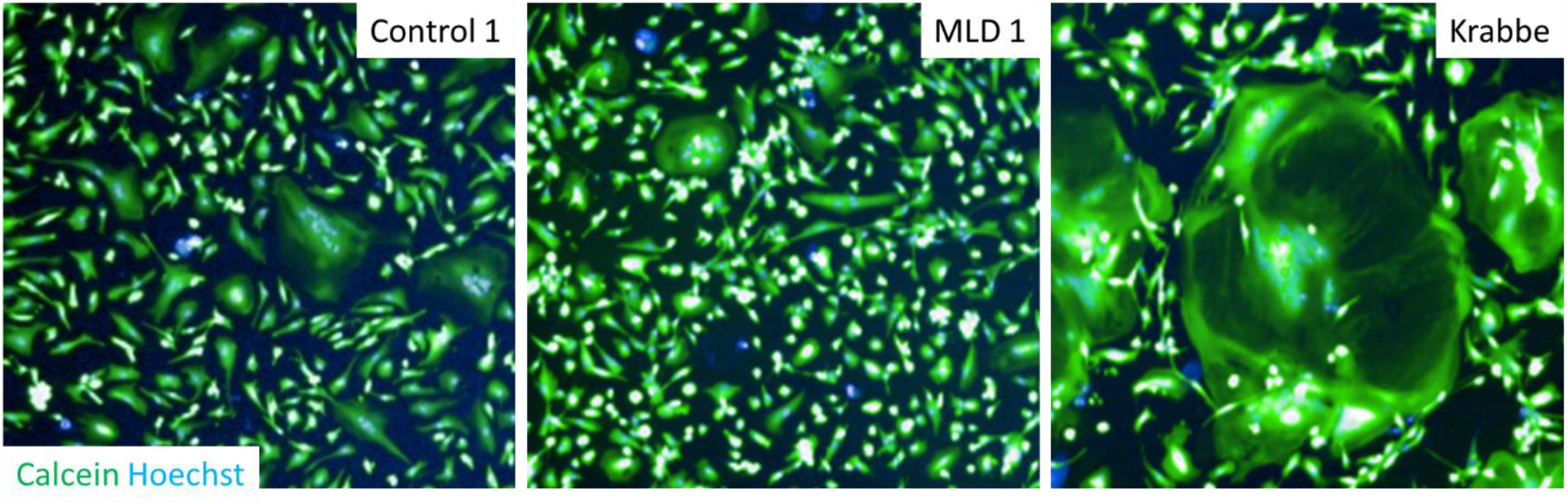
Cropped view of multinucleated cells from Control 1, MLD1, and the Krabbe donor indicated in Figure 1F. The image region highlighted in Figure 2F with white and red arrows has been cropped using the same dimensions to illustrate the difference in multinucleated cells that form in microglia cultures following passaging from the Control 1, MLD1, and Krabbe donor. These have been cropped from the same images shown in Figure 2F. Nuclei are shown in blue, live cells are shown in green.

**Supplementary figure 2.**
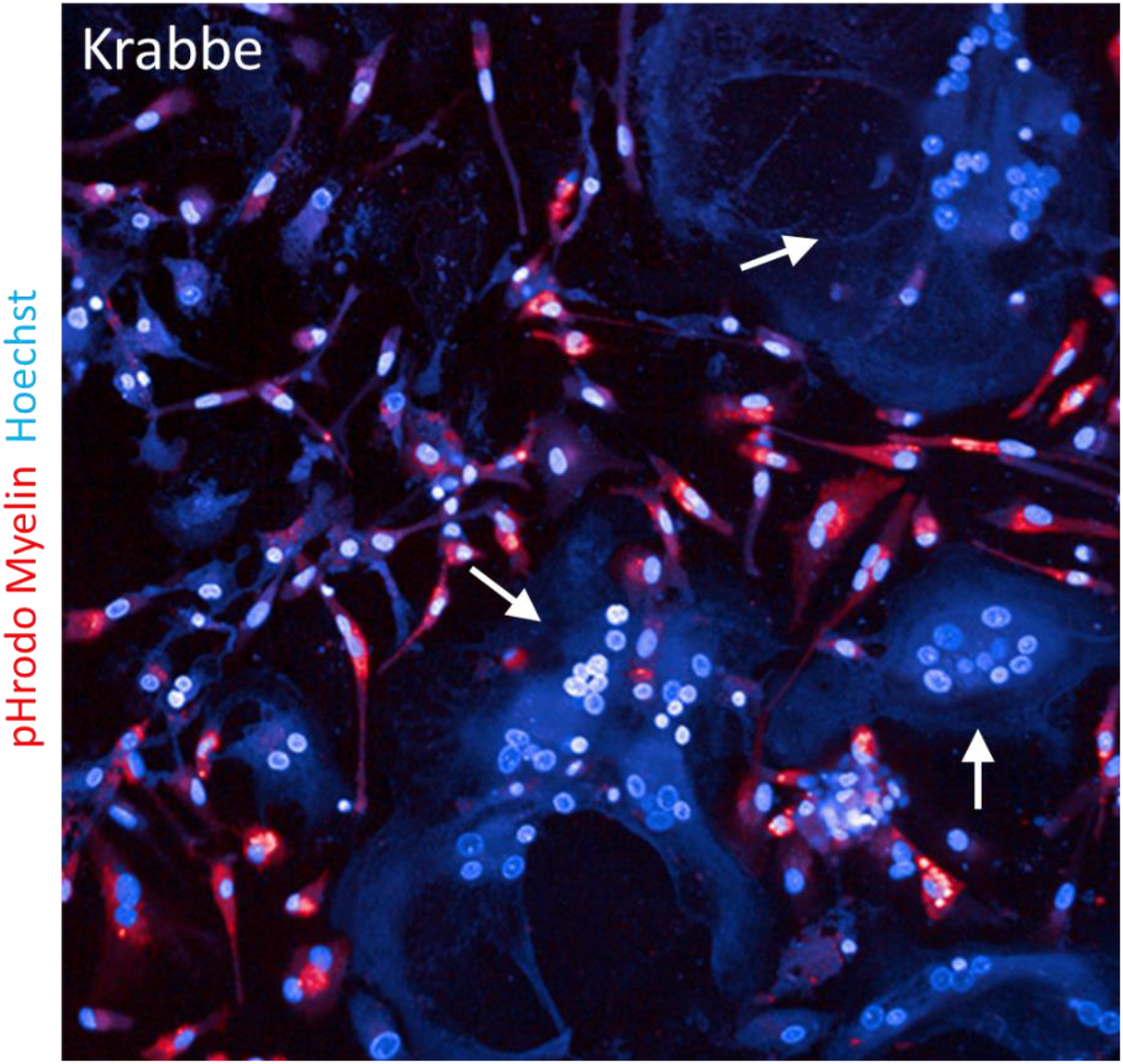
Additional example of impaired phagocytosis of myelin debris in Krabbe multinucleated globoid cells. An additional example from the experiment assessing phagocytosis in Krabbe microglia depicted in figure 4. Microglia from the Krabbe donor were passaged and seeded into a 96 well assay plate for 96 hours. Cells were incubated with 20 µg/mL pHrodo-conjugated myelin debris for 2 hours, counterstained with Hoechst, and then imaged live with a 20x objective. White arrows indicated multinucleated globoid cells that failed to internalize the conjugated substrate. pHrodo myelin is pseudo colored red.

**Supplementary figure 3.**
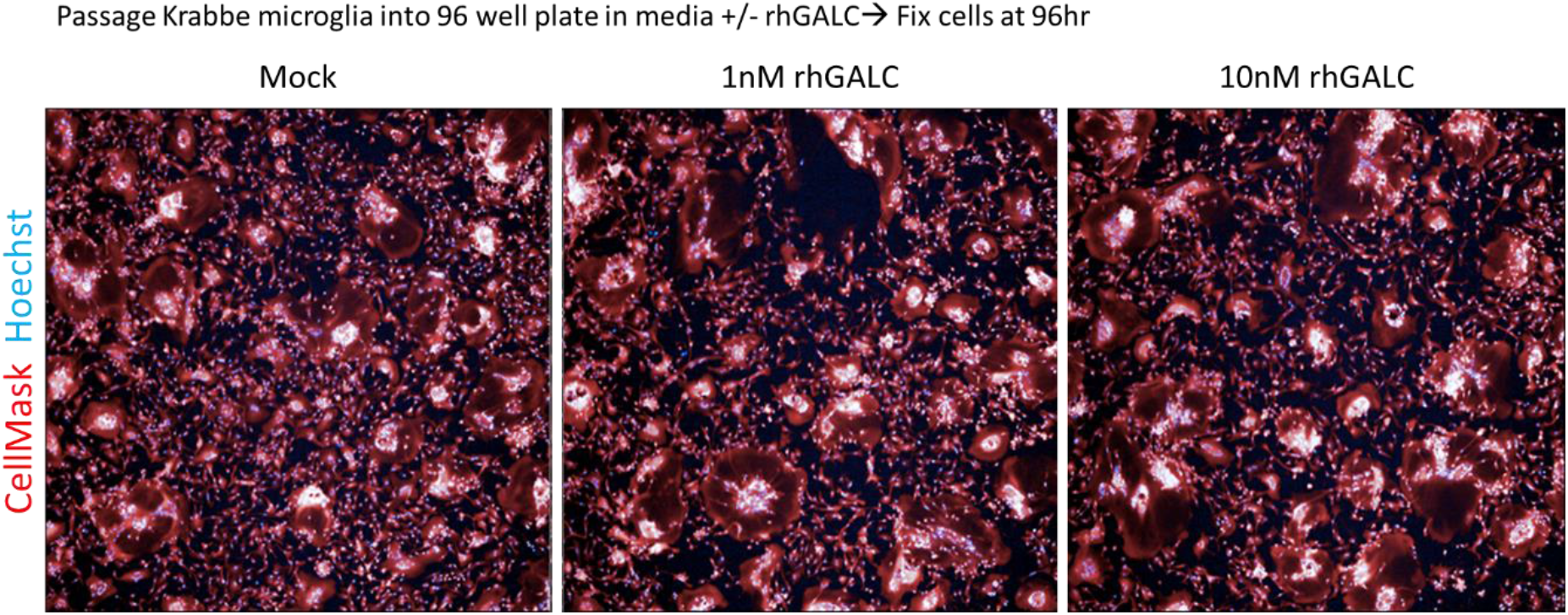
Addition of rhGALC to the culture medium following passaging has no effect on globoid cell formation. Microglia from the Krabbe donor were passaged and seeded into a 96 well assay plate in microglia medium supplemented with 1 nM and 10 nM rhGALC. Cells were cultured for 96 hours prior to fixation. We did not observe a difference in the formation of globoid cells when cultured in the presence of rhGALC protein. Images were obtained with a 5x objective.

## References

1. Hannun YA, Obeid LM. Sphingolipids and their metabolism in physiology and disease. Nat Rev Mol Cell Biol. 2018;19(3):175–91.

2. Iwabuchi K, Nakayama H, Oizumi A, Suga Y, Ogawa H, Takamori K. Role of Ceramide from Glycosphingolipids and Its Metabolites in Immunological and Inflammatory Responses in Humans. Mediumtors Inflamm. 2015;2015:120748.

3. Lingwood CA. Glycosphingolipid functions. Cold Spring Harb Perspect Biol. 2011;3(7).

4. Schnaar RL, Kinoshita T. Glycosphingolipids. In: rd, Varki A, Cummings RD, Esko JD, Stanley P, Hart GW, et al., editors. Essentials of Glycobiology. Cold Spring Harbor (NY)2015. p. 125–35.

5. Sabourdy F, Astudillo L, Colacios C, Dubot P, Mrad M, Segui B, et al. Monogenic neurological disorders of sphingolipid metabolism. Biochim Biophys Acta. 2015;1851(8):1040–51.

6. Schulze H, Sandhoff K. Sphingolipids and lysosomal pathologies. Biochim Biophys Acta. 2014;1841(5):799–810.

7. Kohlschutter A. Lysosomal leukodystrophies: Krabbe disease and metachromatic leukodystrophy. Handb Clin Neurol. 2013;113:1611–8.

8. Kobayashi T, Goto I, Yamanaka T, Suzuki Y, Nakano T, Suzuki K. Infantile and fetal globoid cell leukodystrophy: analysis of galactosylceramide and galactosylsphingosine. Ann Neurol. 1988;24(4):517–22.

9. Orsini JJ, Escolar ML, Wasserstein MP, Caggana M. Krabbe Disease. In: Adam MP, Ardinger HH, Pagon RA, Wallace SE, Bean LJH, Stephens K, et al., editors. GeneReviews((R)). Seattle (WA)1993.

10. Suzuki K. Globoid cell leukodystrophy (Krabbe’s disease): update. J Child Neurol. 2003;18(9):595–603.

11. Zlotogora J, Chakraborty S, Knowlton RG, Wenger DA. Krabbe disease locus mapped to chromosome 14 by genetic linkage. Am J Hum Genet. 1990;47(1):37–44.

12. Suzuki K, Suzuki Y. Globoid cell leucodystrophy (Krabbe’s disease): deficiency of galactocerebroside beta-galactosidase. Proc Natl Acad Sci U S A. 1970;66(2):302–9.

13. Wenger DA, Rafi MA, Luzi P, Datto J, Costantino-Ceccarini E. Krabbe disease: genetic aspects and progress toward therapy. Mol Genet Metab. 2000;70(1):1–9.

14. Svennerholm L, Vanier MT, Mansson JE. Krabbe disease: a galactosylsphingosine (psychosine) lipidosis. J Lipid Res. 1980;21(1):53–64.

15. White AB, Givogri MI, Lopez-Rosas A, Cao H, van Breemen R, Thinakaran G, et al. Psychosine accumulates in membrane microdomains in the brain of krabbe patients, disrupting the raft architecture. J Neurosci. 2009;29(19):6068–77.

16. Won JS, Kim J, Paintlia MK, Singh I, Singh AK. Role of endogenous psychosine accumulation in oligodendrocyte differentiation and survival: implication for Krabbe disease. Brain Res. 2013;1508:44–52.

17. Giri S, Khan M, Rattan R, Singh I, Singh AK. Krabbe disease: psychosine-mediumted activation of phospholipase A2 in oligodendrocyte cell death. J Lipid Res. 2006;47(7):1478–92.

18. Castelvetri LC, Givogri MI, Zhu H, Smith B, Lopez-Rosas A, Qiu X, et al. Axonopathy is a compounding factor in the pathogenesis of Krabbe disease. Acta Neuropathol. 2011;122(1):35–48.

19. Smith BR, Santos MB, Marshall MS, Cantuti-Castelvetri L, Lopez-Rosas A, Li G, et al. Neuronal inclusions of alpha-synuclein contribute to the pathogenesis of Krabbe disease. J Pathol. 2014;232(5):509–21.

20. O’Sullivan C, Dev KK. Galactosylsphingosine (psychosine)-induced demyelination is attenuated by sphingosine 1-phosphate signalling. J Cell Sci. 2015;128(21):3878–87.

21. Misslin C, Velasco-Estevez M, Albert M, O’Sullivan SA, Dev KK. Phospholipase A2 is involved in galactosylsphingosine-induced astrocyte toxicity, neuronal damage and demyelination. PLoS One. 2017;12(11):e0187217.

22. Claycomb KI, Winokur PN, Johnson KM, Nicaise AM, Giampetruzzi AW, Sacino AV, et al. Aberrant production of tenascin-C in globoid cell leukodystrophy alters psychosine-induced microglial functions. J Neuropathol Exp Neurol. 2014;73(10):964–74.

23. Ijichi K, Brown GD, Moore CS, Lee JP, Winokur PN, Pagarigan R, et al. MMP-3 mediumtes psychosine-induced globoid cell formation: implications for leukodystrophy pathology. Glia. 2013;61(5):765–77.

24. Suzuki K. Twenty five years of the “psychosine hypothesis”: a personal perspective of its history and present status. Neurochem Res. 1998;23(3):251–9.

25. Suzuki K. Ultrastructural study of experimental globoid cells. J Neuropathol Exp Neurol. 1971;30(1):145–6.

26. Itoh M, Hayashi M, Fujioka Y, Nagashima K, Morimatsu Y, Matsuyama H. Immunohistological study of globoid cell leukodystrophy. Brain Dev. 2002;24(5):284–90.

27. Borda JT, Alvarez X, Mohan M, Ratterree MS, Phillippi-Falkenstein K, Lackner AA, et al. Clinical and immunopathologic alterations in rhesus macaques affected with globoid cell leukodystrophy. Am J Pathol. 2008;172(1):98–111.

28. McInnes A, Rennick DM. Interleukin 4 induces cultured monocytes/macrophages to form giant multinucleated cells. J Exp Med. 1988;167(2):598–611.

29. McNally AK, Anderson JM. Multinucleated giant cell formation exhibits features of phagocytosis with participation of the endoplasmic reticulum. Exp Mol Pathol. 2005;79(2):126–35.

30. Brodbeck WG, Anderson JM. Giant cell formation and function. Curr Opin Hematol. 2009;16(1):53–7.

31. Nicaise AM, Bongarzone ER, Crocker SJ. A microglial hypothesis of globoid cell leukodystrophy pathology. J Neurosci Res. 2016;94(11):1049–61.

32. Potter GB, Santos M, Davisson MT, Rowitch DH, Marks DL, Bongarzone ER, et al. Missense mutation in mouse GALC mimics human gene defect and offers new insights into Krabbe disease. Hum Mol Genet. 2013;22(17):3397–414.

33. Gomez-Ospina N. Arylsulfatase A Deficiency. In: Adam MP, Ardinger HH, Pagon RA, Wallace SE, Bean LJH, Stephens K, et al., editors. GeneReviews((R)). Seattle (WA)1993.

34. Biffi A, Lucchini G, Rovelli A, Sessa M. Metachromatic leukodystrophy: an overview of current and prospective treatments. Bone Marrow Transplant. 2008;42 Suppl 2:S2–6.

35. Gieselmann V, Krageloh-Mann I. Metachromatic leukodystrophy--an update. Neuropediatrics. 2010;41(1):1–6.

36. Gieselmann V, Polten A, Kreysing J, von Figura K. Molecular genetics of metachromatic leukodystrophy. J Inherit Metab Dis. 1994;17(4):500–9.

37. Gieselmann V. Metachromatic leukodystrophy: genetics, pathogenesis and therapeutic options. Acta Paediatr. 2008;97(457):15–21.

38. Hess B, Saftig P, Hartmann D, Coenen R, Lullmann-Rauch R, Goebel HH, et al. Phenotype of arylsulfatase A-deficient mice: relationship to human metachromatic leukodystrophy. Proc Natl Acad Sci U S A. 1996;93(25):14821–6.

39. Bergner CG, van der Meer F, Winkler A, Wrzos C, Turkmen M, Valizada E, et al. Microglia damage precedes major myelin breakdown in X-linked adrenoleukodystrophy and metachromatic leukodystrophy. Glia. 2019;67(6):1196–209.

40. Biffi A, Montini E, Lorioli L, Cesani M, Fumagalli F, Plati T, et al. Lentiviral hematopoietic stem cell gene therapy benefits metachromatic leukodystrophy. Science. 2013;341(6148):1233158.

41. Biffi A, Capotondo A, Fasano S, del Carro U, Marchesini S, Azuma H, et al. Gene therapy of metachromatic leukodystrophy reverses neurological damage and deficits in mice. J Clin Invest. 2006;116(11):3070–82.

42. Allewelt H, Taskindoust M, Troy J, Page K, Wood S, Parikh S, et al. Long-Term Functional Outcomes after Hematopoietic Stem Cell Transplant for Early Infantile Krabbe Disease. Biol Blood Marrow Transplant. 2018;24(11):2233–8.

43. Suzuki K. Biochemical pathogenesis of genetic leukodystrophies: comparison of metachromatic leukodystrophy and globoid cell leukodystrophy (Krabbe’s disease). Neuropediatrics. 1984;15 Suppl:32–6.

44. Takahashi K, Tanabe K, Ohnuki M, Narita M, Ichisaka T, Tomoda K, et al. Induction of pluripotent stem cells from adult human fibroblasts by defined factors. Cell. 2007;131(5):861–72.

45. Mertens J, Marchetto MC, Bardy C, Gage FH. Evaluating cell reprogramming, differentiation and conversion technologies in neuroscience. Nat Rev Neurosci. 2016;17(7):424–37.

46. Abud EM, Ramirez RN, Martinez ES, Healy LM, Nguyen CHH, Newman SA, et al. iPSC-Derived Human Microglia-like Cells to Study Neurological Diseases. Neuron. 2017;94(2):278–93 e9.

47. Muffat J, Li Y, Yuan B, Mitalipova M, Omer A, Corcoran S, et al. Efficient derivation of microglia-like cells from human pluripotent stem cells. Nat Med. 2016;22(11):1358–67.

48. Douvaras P, Sun B, Wang M, Kruglikov I, Lallos G, Zimmer M, et al. Directed Differentiation of Human Pluripotent Stem Cells to Microglia. Stem Cell Reports. 2017;8(6):1516–24.

49. Brownjohn PW, Smith J, Solanki R, Lohmann E, Houlden H, Hardy J, et al. Functional Studies of Missense TREM2 Mutations in Human Stem Cell-Derived Microglia. Stem Cell Reports. 2018;10(4):1294–307.

50. Haenseler W, Sansom SN, Buchrieser J, Newey SE, Moore CS, Nicholls FJ, et al. A Highly Efficient Human Pluripotent Stem Cell Microglia Model Displays a Neuronal-Co-culture-Specific Expression Profile and Inflammatory Response. Stem Cell Reports. 2017;8(6):1727–42.

51. van Wilgenburg B, Browne C, Vowles J, Cowley SA. Efficient, long term production of monocyte-derived macrophages from human pluripotent stem cells under partly-defined and fully-defined conditions. PLoS One. 2013;8(8):e71098.

52. Scott-Hewitt NJ, Folts CJ, Hogestyn JM, Piester G, Mayer-Proschel M, Noble MD. Heterozygote galactocerebrosidase (GALC) mutants have reduced remyelination and impaired myelin debris clearance following demyelinating injury. Hum Mol Genet. 2017;26(15):2825–37.

53. Tcw J, Wang M, Pimenova AA, Bowles KR, Hartley BJ, Lacin E, et al. An Efficient Platform for Astrocyte Differentiation from Human Induced Pluripotent Stem Cells. Stem Cell Reports. 2017;9(2):600–14.

54. Binder F, Hayakawa M, Choo MK, Sano Y, Park JM. Interleukin-4-induced beta-catenin regulates the conversion of macrophages to multinucleated giant cells. Mol Immunol. 2013;54(2):157–63.

55. Takata K, Kozaki T, Lee CZW, Thion MS, Otsuka M, Lim S, et al. Induced-Pluripotent-Stem-Cell-Derived Primitive Macrophages Provide a Platform for Modeling Tissue-Resident Macrophage Differentiation and Function. Immunity. 2017;47(1):183–98 e6.

56. Kondo Y, Adams JM, Vanier MT, Duncan ID. Macrophages counteract demyelination in a mouse model of globoid cell leukodystrophy. J Neurosci. 2011;31(10):3610–24.

57. Neumann H, Kotter MR, Franklin RJ. Debris clearance by microglia: an essential link between degeneration and regeneration. Brain. 2009;132(Pt 2):288–95.

58. Safaiyan S, Kannaiyan N, Snaidero N, Brioschi S, Biber K, Yona S, et al. Age-related myelin degradation burdens the clearance function of microglia during aging. Nat Neurosci. 2016;19(8):995–8.

59. LeVine SM, Brown DC. IL-6 and TNFalpha expression in brains of twitcher, quaking and normal mice. J Neuroimmunol. 1997;73(1-2):47–56.

60. Giri S, Jatana M, Rattan R, Won JS, Singh I, Singh AK. Galactosylsphingosine (psychosine)-induced expression of cytokine-mediumted inducible nitric oxide synthases via AP-1 and C/EBP: implications for Krabbe disease. FASEB J. 2002;16(7):661–72.

61. Potter GB, Petryniak MA. Neuroimmune mechanisms in Krabbe’s disease. J Neurosci Res. 2016;94(11):1341–8.

62. Snook ER, Fisher-Perkins JM, Sansing HA, Lee KM, Alvarez X, MacLean AG, et al. Innate immune activation in the pathogenesis of a murine model of globoid cell leukodystrophy. Am J Pathol. 2014;184(2):382–96.

63. Pedchenko TV, LeVine SM. IL-6 deficiency causes enhanced pathology in Twitcher (globoid cell leukodystrophy) mice. Exp Neurol. 1999;158(2):459–68.

64. Moreno JL, Mikhailenko I, Tondravi MM, Keegan AD. IL-4 promotes the formation of multinucleated giant cells from macrophage precursors by a STAT6-dependent, homotypic mechanism: contribution of E-cadherin. J Leukoc Biol. 2007;82(6):1542–53.

65. Dadsetan M, Jones JA, Hiltner A, Anderson JM. Surface chemistry mediumtes adhesive structure, cytoskeletal organization, and fusion of macrophages. J Biomed Mater Res A. 2004;71(3):439–48.

66. Deane JE, Graham SC, Kim NN, Stein PE, McNair R, Cachon-Gonzalez MB, et al. Insights into Krabbe disease from structures of galactocerebrosidase. Proc Natl Acad Sci U S A. 2011;108(37):15169–73.

67. Marques AR, Willems LI, Herrera Moro D, Florea BI, Scheij S, Ottenhoff R, et al. A Specific Activity-Based Probe to Monitor Family GH59 Galactosylceramidase, the Enzyme Deficient in Krabbe Disease. Chembiochem. 2017;18(4):402–12.

68. Vignery A. Macrophage fusion: are somatic and cancer cells possible partners? Trends Cell Biol. 2005;15(4):188–93.

69. Vignery A. Macrophage fusion: the making of osteoclasts and giant cells. J Exp Med. 2005;202(3):337–40.

70. Fendrick SE, Xue QS, Streit WJ. Formation of multinucleated giant cells and microglial degeneration in rats expressing a mutant Cu/Zn superoxide dismutase gene. J Neuroinflammation. 2007;4:9.

71. Budka H. Multinucleated giant cells in brain: a hallmark of the acquired immune deficiency syndrome (AIDS). Acta Neuropathol. 1986;69(3-4):253–8.

72. Ferrer I, Boada Rovira M, Sanchez Guerra ML, Rey MJ, Costa-Jussa F. Neuropathology and pathogenesis of encephalitis following amyloid-beta immunization in Alzheimer’s disease. Brain Pathol. 2004;14(1):11–20.

